# Genetic background influences survival of infections with *Salmonella enterica* serovar Typhimurium in the Collaborative Cross

**DOI:** 10.1101/2022.02.07.479341

**Authors:** Kristin Scoggin, Rachel Lynch, Jyotsana Gupta, Aravindh Nagarajan, Max Sheffield, Ahmed Elsaadi, Christopher Bowden, Manuchehr Aminian, Amy Peterson, L. Garry Adams, Michael Kirby, David W. Threadgill, Helene Andrews-Polymenis

**Author notes:** Corresponding authors: (DWT), (HAP), (MK). Department of Genetics, University of North Carolina, Chapel Hill, North Carolina, United States of America (Rachel Lynch). PerkinElmer Inc., Hopkinton, Massachusetts, United States of America (Jyotsana Gupta). Department of Mathematics and Statistics, California State Polytechnic University, Pomona, California, United States of America (Manuchehr Aminian).

## Abstract

*Salmonella* infections typically cause self-limiting gastroenteritis, but in some individuals these bacteria can spread systemically and cause disseminated disease. *Salmonella* Typhimurium (STm), which causes severe systemic disease in most inbred mice, has been used as a model for disseminated disease. To screen for new infection phenotypes across a range of host genetics, we orally infected 32 Collaborative Cross (CC) mouse strains with STm and monitored their disease progression for seven days by telemetry. Our data revealed a broad range of phenotypes across CC strains in many parameters including survival, bacterial colonization, tissue damage, complete blood counts (CBC), and serum cytokines. Eighteen CC strains survived to day 7, while fourteen susceptible strains succumbed to infection before day 7. Several CC strains had sex differences in survival and colonization. Surviving strains had lower pre-infection baseline temperatures and were less active during their daily active period. Core body temperature disruptions were detected earlier after STm infection than activity disruptions, making temperature a better detector of illness. All CC strains had STm in spleen and liver, but susceptible strains were more highly colonized. Tissue damage was weakly negatively correlated to survival. We identified loci associated with survival on Chromosomes (Chr) 1, 2, 4, 7. Polymorphisms in *Ncf2* and *Slc11a1*, known to reduce survival in mice after STm infections, are located in the Chr 1 interval, and the Chr 7 association overlaps with a previously identified QTL peak called *Ses2*. We identified two new genetic regions on Chr 2 and 4 associated with susceptibility to STm infection. Our data reveal the diversity of responses to STm infection across a range of host genetics and identified new candidate regions for survival of STm infection.

**Author Summary:** *Salmonella* Typhimurium (STm) infections typically cause self-limiting diarrheal symptoms, but in some individuals, the bacteria can spread throughout the body and cause life-threatening infection. We used a population of genetically different mice (Collaborative Cross) to identify their range of responses to STm infection. We identified a broad range of outcomes across these different mice, including a group of mice susceptible to lethal infection and a group that survived our 7 day study. We found that mice that survived STm infection had a cooler core body temperature before infection than susceptible mice, while remaining active. Thus, body temperature, rather than activity, appears to be a better predictor of poor outcomes after STm infection. We identified several regions of the mouse genome that are associated with outcome after STm infection. One of these regions, mouse Chromosome (Chr) 1 has genes that are already known to influence susceptibility to STm infection. Two other regions that we identified to influence survival after STm infection, located on mouse Chr 2 and 4, are novel and contain numerous genes of interest that may be linked to susceptibility. Our work defines the utility of exploring how host genetic diversity influences infection outcomes with bacterial pathogens.

## Introduction

Salmonellae are Gram negative facultative intracellular bacteria that cause self-limiting gastroenteritis, typhoid fever, or sepsis in human beings (1–3). Non-typhoidal *Salmonella* (NTS) infections are estimated to cause 93.8 million human cases per year, resulting in 155,000 deaths worldwide (4). While NTS infection is typically limited to the intestine and causes diarrheal disease, NTS can also cause more severe disseminated disease, depending on the *Salmonella* serotype and host factors such as age, immune status, and genetics (5). Worldwide, 535,000 human cases of NTS disseminated disease, either *Salmonella* bacteremia or invasive non-typhoidal salmonellosis, occur annually and cause 77,500 deaths (6).

Although antibiotics are not generally used to treat patients with NTS gastroenteritis, they are used to treat patients with NTS bacteremia and invasive non-typhoidal salmonellosis (iNTS). Thus, a concern of increasing importance is the development of multi-drug resistant strains of *Salmonella* that cause 212,500 human cases per year in the US alone, with 16% of all NTS cases resistant to at least one antibiotic as of 2017 (7). Exploitation of differences in host immunity may help identify new interventions that could become treatment alternatives to antibiotics (8–10).

*Salmonella enterica* serotype Typhimurium (STm) is a serotype of non-typhoidal *Salmonella* that causes murine typhoid resembling severe invasive human infections. Some mouse strains are particularly susceptible (C57BL/6J and BALB/cJ), while others are able to resist severe systemic disease (CBA/J, 129/SvJ, and A/J) (11–13). Previous studies have largely attributed differential susceptibility to a mutation in solute carrier family 11 member 1 (*Slc11a1*) encoding a metal cation pump in macrophages that reduces intracellular growth of bacteria (2,14–17). Natural variation in other genes has also been shown to play a role in differential survival after STm infection, including neutrophil cytosolic factor 2 (*Ncf2*) and toll-like receptor 4 (*Tlr4*). *Ncf2* encodes a component of the NADPH complex in neutrophils that generates superoxide and is vital to neutrophil’s killing ability (18–20). Toll-like receptor 4 (*Tlr4*) recognizes lipopolysaccharide (LPS) and is a key component of innate immune response activation and cytokine production (21,22).

To identify new host genetic mechanisms that could influence differential susceptibility to STm infections, we used the Collaborative Cross (CC) mouse genetic reference population (23–26). The CC has greater variability across all parameters than traditional strains, including immune phenotypes, because of its genetic diversity (27–29). This population has been used to identify novel genetic variants and discover new phenotypes in mice, such as tolerance (high pathogen load with minimal disease) (30–32) for infectious agents including Influenza A, Ebola, *Mycobacterium tuberculosis*, Theiler’s murine encephalomyelitis virus, SARS-Corona virus, and West Nile virus (33–39).

To identify new mechanisms underlying diverse host responses to STm infections in mice, we infected CC strains orally and monitored infected animals for one-week post-infection using telemetry to track temperature and activity levels. At necropsy, organs and blood were collected to analyze colonization, histopathology, complete blood counts (CBC), and cytokine levels. Strains that survived to day 7 had lower uninfected baseline body temperatures and were less active during their active period pre-infection. Temperature patterns were disrupted earlier than activity patterns, suggesting that temperature is a more sensitive measurement for illness. All parameters were strain dependent, but in general, mice were colonized in systemic organs independent of survival time, spleen damage was most correlated to survival, and cytokines and CBC parameters were not predictive of survival. Several CC strains showed sex differences across survival, weight change, and bacterial burden, but one strain, CC027/GeniUnc, showed stark sex-dependence across all parameters. Quantitative trait locus (QTL) analysis for survival time revealed three significant and two suggestive associations, confirming known loci and detecting previously unknown variants providing new candidate genes not previously associated with STm. Furthermore, despite having susceptible alleles in both *Slc11a1* and *Ncf2* that are linked to early death from *Salmonella* infection, CC045/GeniUnc mice were able to survive 7 days with a high bacterial load and exhibited tolerance to systemic *Salmonella* infection.

## Results

### Response to oral *Salmonella* Typhimurium infection is variable across CC strains

We infected thirty-two Collaborative Cross (CC) strains (3 male and 3 female) with *S.* Typhimurium (STm) ATCC14028 and monitored them for up to one week to identify variable susceptibility to STm infection. Due to their well-characterized response to STm infection, C57BL/6J (B6) mice, which develop systemic murine typhoid, and CBA/J (CBA) mice, which are resistant to systemic infection, were used as infection controls. At the time of necropsy, sections of ileum, cecum, colon, liver, and spleen were collected to determine bacterial load. We focused our analysis on the systemic phase of infection. In comparison to previous studies that injected STm intravenously into CC mice, our STm oral infections produced a much wider range of colonization in critical organs (Fig. 1). Using sex-combined median data, the correlation between liver and spleen colonization with survival time was R = -0.62 and -0.63, respectively (Supp. Fig. 1).

**Figure 1:**
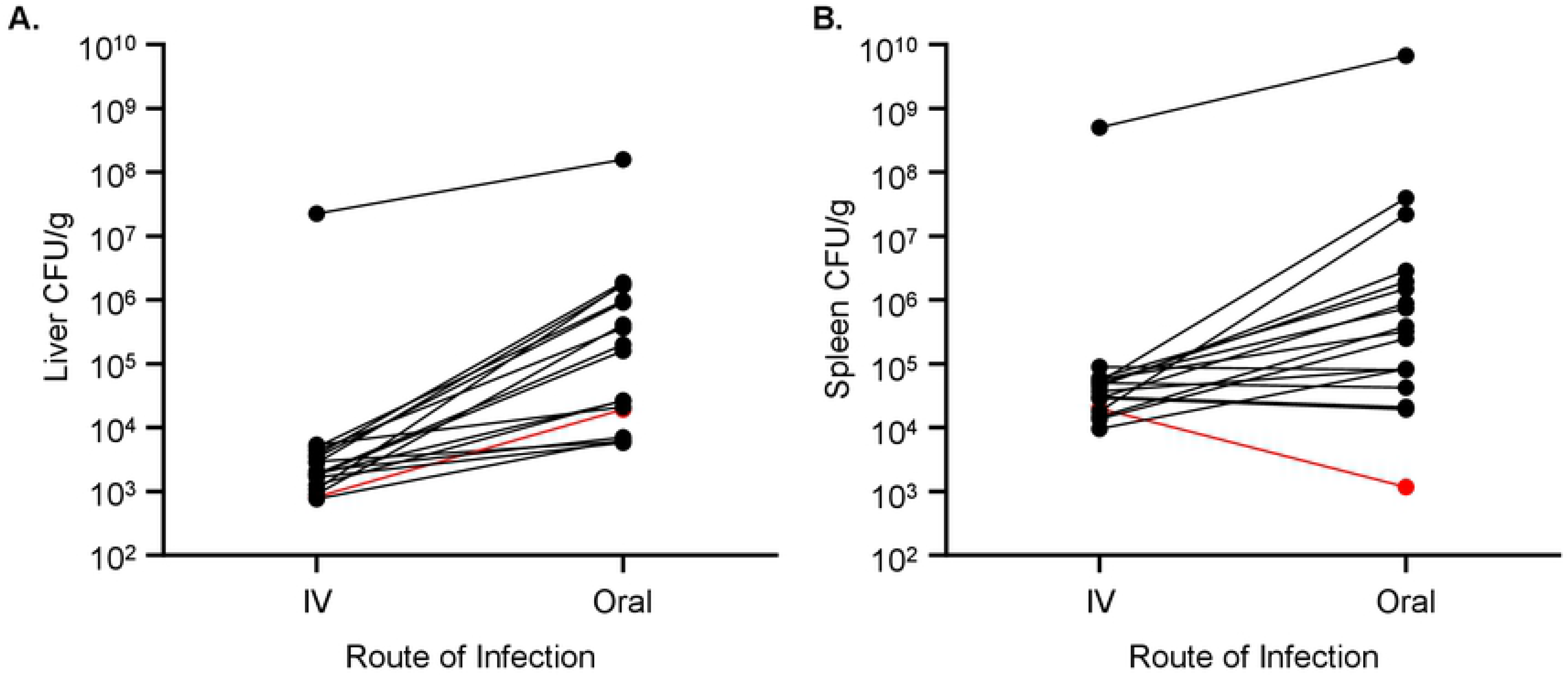
CC strains infected orally with STm are more highly and variably colonized than after IV infection. A. Liver and B. spleen colonization (median) for 18 CC strains orally infected in this project and infected intravenously in the previous work by Zhang, J. et al. 2018. Strains are connected by lines, and the red line represents CC051.

All B6 mice met the euthanasia criteria between day 3 and 6 post-infection with a median at day 4 and were classified as highly susceptible (Fig. 2A). B6 mice lost a median of 20.73% of their body weight during infection (Fig. 2B). CBA mice survived to day 7 and were classified as resistant (Fig. 2A) with a median loss of 7.92% of their body weight during infection (Fig. 2B). B6 mice were highly colonized in the liver and spleen with medians of 2.56 x10^6^ and 1.21 x10^7^ CFU/g, respectively (Fig. 2C, D), while CBA mice were colonized at least two orders of magnitude lower with medians of 5.45 x10^3^ and 1.84 x10^4^ CFU/g, respectively (Fig. 2C, D).

**Figure 2:**
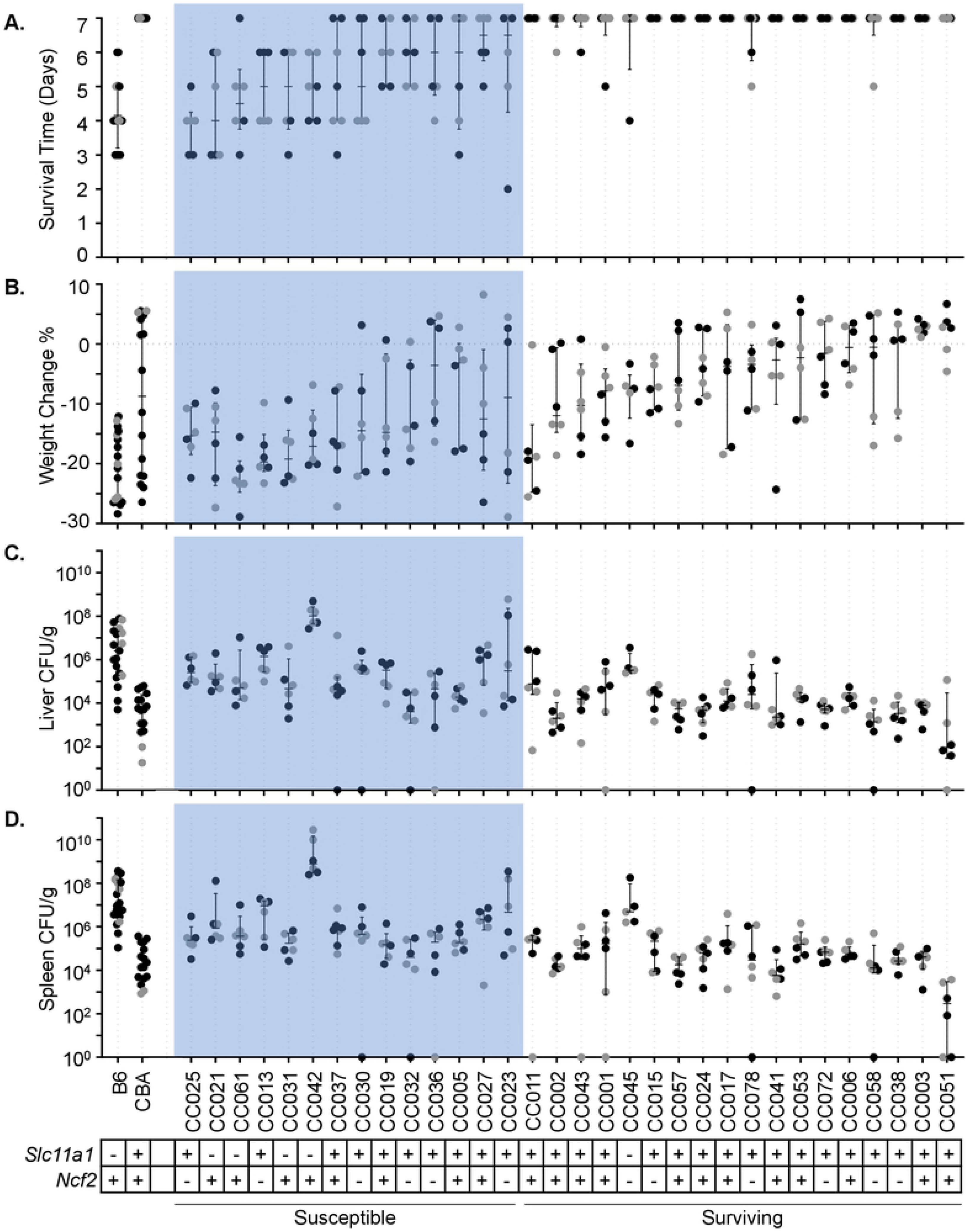
CC strains have variable responses to STm infection. A. Survival time and B. percent weight change after infection with STm. Strains are in ascending order of survival and weight change if survival was equal between strains. Bacterial burden in C. liver and D. spleen. Dots represent individual mice; black dots represent males and grey dots represent females. Median and interquartile range are shown by lines for each strain. *Slc11a1* and *Ncf2* status shown by +/-. 32 CC strains and 2 controls (B6 and CBA) represented.

Of the 32 CC strains we infected, 14 were categorized as susceptible (3 or more mice out of 6 infected met euthanasia criteria prior to day 7 post-infection) and 18 strains were called resistant or tolerant (Fig. 2A, susceptible in shaded region). Susceptible strains that suffered earlier mortality also had greater weight loss (Fig. 2B) with a correlation of R = 0.67 (Supp. Fig. 2). Susceptible strains had a least 10-fold lower bacterial burden in liver than B6 control mice, with the exception of CC013, CC027, and CC042 (Fig. 2C). CC013 and CC027 had comparable colonization in liver to B6, and CC042 had 100-fold higher with a median colonization of 1.02 x10^8^ CFU/g (Fig. 2C), consistent with previous reports (40,41). In spleen, all strains had at least a 10-fold lower bacterial burden than B6 mice, with the exception of CC042 (Fig. 2D), which were colonized 100-fold higher than B6 mice (Fig. 2D). The high colonization of CC042 has been previously attributed to a *de novo* mutation in integrin alpha (*Itgal*) (41). Susceptible strains were also colonized at least 10-fold higher than CBA control mice, with the exception of CC032 (Fig. 2C, D). CC032 mice were colonized in both liver and spleen similarly to CBA with a median of 4.11 x10^3^ and 3.96 x10^4^ CFU/g, respectively, yet succumbed to infection before day 7 (Fig. 2C, 2D).

An additional 18 strains were categorized as resistant or tolerant. Resistance was defined as surviving infection by preventing colonization or by clearing the bacteria. Tolerance was defined as surviving infection while maintaining high bacterial burden. Both resistant and tolerant strains had </= 2 mice meeting euthanasia criteria prior to day 7. Of these 18 strains, all 6 mice lived to day 7 in 12 strains (Fig. 2A). Strains that survived longer lost less weight than those that did not with a correlation of R = 0.71 across all strains (Fig. 2B and Supp. Fig. 1). However, CC011 lost substantially more weight than expected for a surviving strain. All the resistant and tolerant strains, with two exceptions (CC045 and CC051), were equally colonized across systemic organs, thus making these phenotypes difficult to differentiate.

One strain – CC051 – had 100-fold lower bacterial burden than CBA in both liver and spleen with medians of 9.30 x10^1^ and 2.92 x10^2^ CFU/g, respectively, and it therefore met our criteria for resistance (Fig. 2C, D). Another strain – CC045 – had a 100-fold higher bacterial burden than CBA mice with 3.31 x10^5^ and 4.83 x10^6^ CFU/g, respectively, making it the strongest candidate for tolerance (Fig. 2C, D). Of the remaining 16 surviving strains, eight were colonized in the liver to the same level as CBA mice, while eight additional strains were colonized in the liver 10-fold higher than CBA mice (Fig. 2C).

### Sex-dependent variation in susceptibility is strain dependent

Sex has been shown to be an influencing factor in many phenotypes, including response to infection (28,42–44). Across the CC population that we tested, sex did not significantly influence survival time (*P* = 0.8149), weight change post-infection (*P* = 0.9764), spleen CFU (*P* = 0.7289), or liver CFU (*P* = 0.5571). However, several individual strains did display sex-dependent variation. CC013 and CC030 males survived longer than females, and CC042 and CC027 females survived longer than males (Fig. 3A). CC015, CC027, and CC072 females lost less weight than males and CC057 males lost less than females (Fig. 3B). CC027 was the only strain where weight loss influenced survival in a sex-dependent manner.

**Figure 3:**
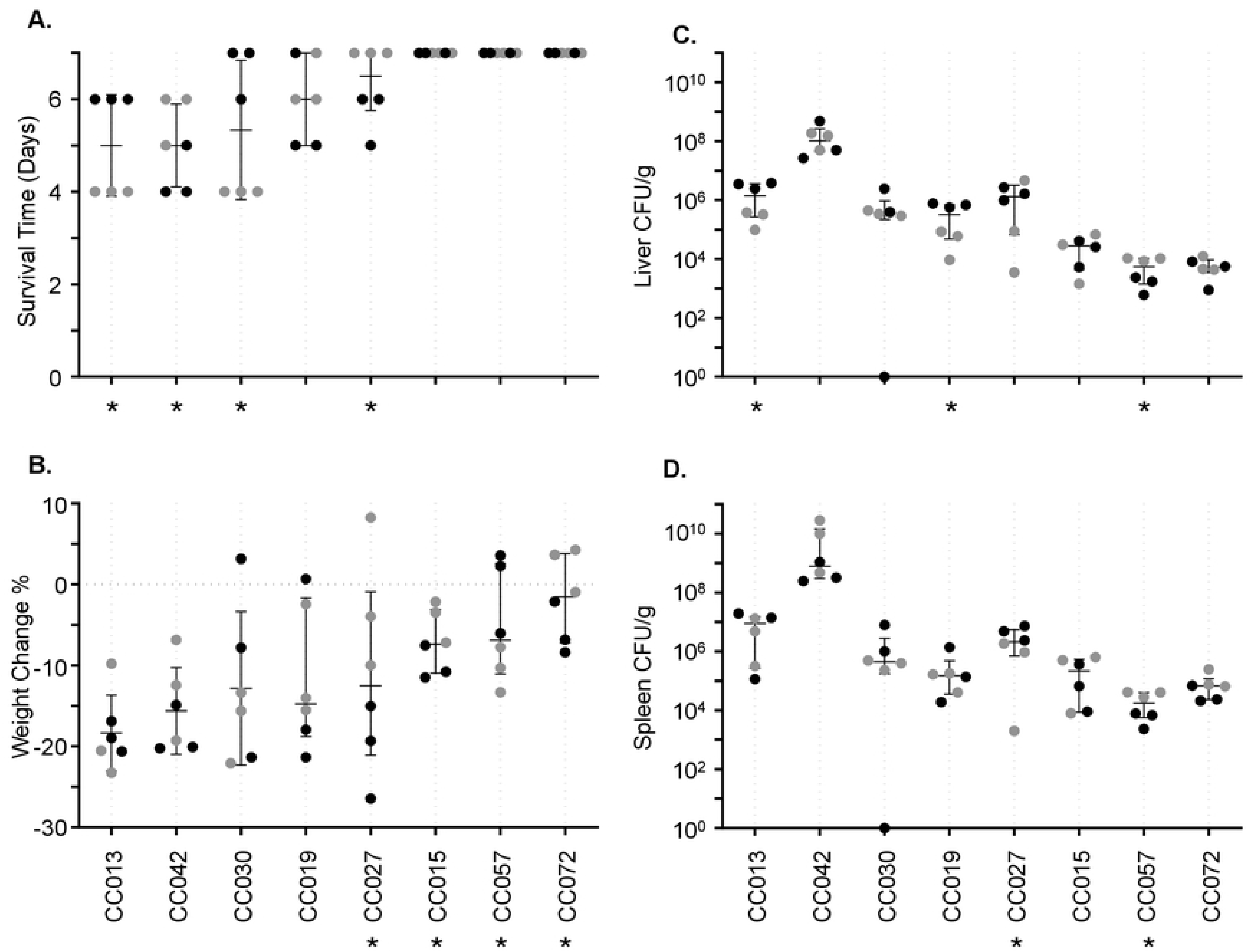
Sex differences in survival after STm infection. A. Survival time and B. percent weight change after infection with STm. Strains are in ascending order of survival and weight change if survival was equal between strains. Bacterial burden in C. liver and D. spleen. Dots represent individual mice; black dots represent males and grey dots represent females. Median and interquartile range are shown by lines for each strain. Eight CC strains are represented. (* P <0.05).

CC013, CC019, and CC027 males had higher bacterial burdens than females, while CC057 females were more highly colonized than males (Fig. 3C, D). CC013 and CC019 had different liver colonization (Fig. 3C), while CC027 had different spleen colonization (Fig. 3D), and CC057 had differences in both organs (Fig. 3C, D). Just as with weight loss, CC027 bacterial burden influenced survival time in a sex-dependent manner. CC027 females survived to day 7, had a lower bacterial burden, and lost less weight than males. As such, CC027 males are susceptible to STm infection, while females are tolerant or resistant.

### Surviving mice have lower baseline body temperatures and stay on baseline circadian pattern for activity longer

Implantable telemetry devices tracked body temperature and activity levels at one-minute intervals both before and throughout infections. Prior to infection, each strain had unique diurnal pattern of baseline temperature and activity prior to infection (Supp. Fig. 3). When strains were grouped by survival outcome, differences between groups became apparent (Fig. 4). Resistant/tolerant mice that survived the infection had a baseline pre-infection median minimum temperature of 36.32 °C during their resting period compared to susceptible non-surviving mice with a pre-infection median minimum temperature of 36.67 °C at rest (Fig. 4A, *P* = <0.0001). The pre-infection median overall temperature for surviving versus susceptible mice was 36.61 °C and 36.95 °C, respectively (Fig. 4A, *P* = <0.0001). This significant difference was also found in maximum temperature reached during the active periods with medians of 37.08 °C and 37.35 °C, for surviving versus non-surviving mice respectively (Fig. 4A, *P* = 0.0025). Thus, surviving strains had a statistically significantly lower pre-infection baseline body temperature relative to susceptible strains.

**Figure 4:**
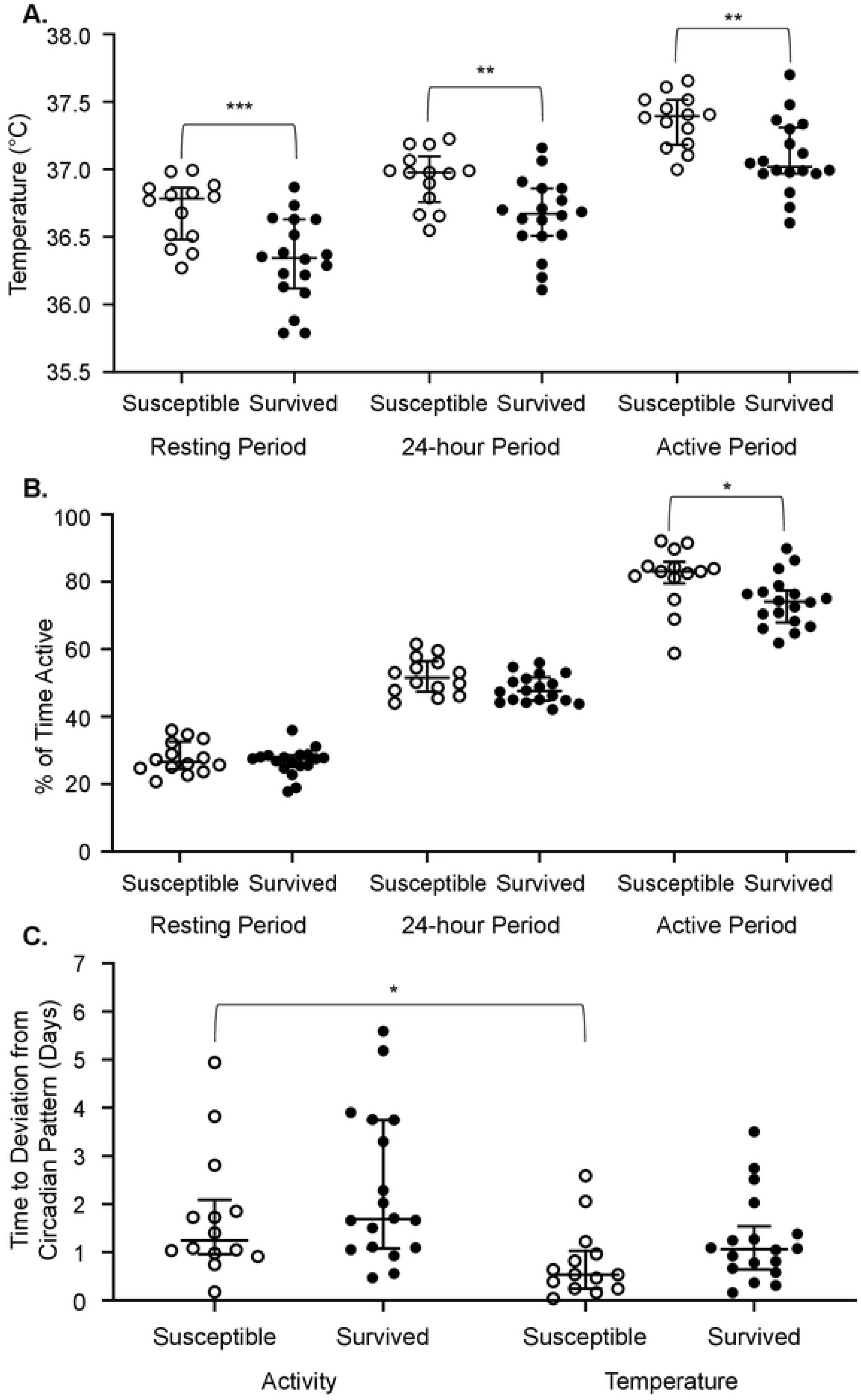
At pre-infection baseline, susceptible strains have higher body temperatures, are more active during normal rest period, and maintain activity patterns longer than temperature patterns than surviving strains. Resting period, 24-hour period, and active period strain medians shown for A. temperature and B. percent of time active. C. Time to deviation from baseline activity and temperature post-infection (days). Strains are grouped by survival status. Median overall values and interquartile range are indicated. (* P <0.05, ** P <0.01, *** P <0.001).

Pre-infection activity measurements were analyzed by what fraction of the time the mouse was active, with a higher fraction suggesting more activity. Before infection, all mice had maximal activity during their active period at night. In pre-infection periods where mice were minimally active (day), both susceptible strains and surviving strains were active for equal amounts of time (Fig. 4B). Considering an entire 24-hour period prior to infection, there was no statistically significant difference in activity between susceptible and surviving strains (Fig. 4B). During the most active periods for the mice, strains susceptible to STm infection spent significantly more time active than strains that survived STm infection (0.83 versus 0.74, Fig. 4B, *P* = 0.0139). Thus, at pre-infection baseline, surviving strains appeared to spend more time at rest during the normal waking period of the mouse (night).

Post-infection telemetry data were analyzed explicitly incorporating assumptions about oscillatory behavior to model when mice deviated from their circadian pattern of body temperature and activity after infection with STm (time to “off-pattern”) (45). For temperature measurements, surviving and susceptible strains did not differ significantly for their time to circadian pattern disruption after infection (Fig. 4C, *P* = 0.3961). Susceptible and surviving strains also did not differ significantly for time to off pattern for activity (Fig. 4C, *P* = >0.9999). However, susceptible strains did stay on their normal activity pattern significantly longer than they stayed on their normal temperature pattern with medians of 1789 minutes (1.24 days) and 770.75 minutes (0.54 days) (Fig. 4C, *P* = 0.0397). Surviving strains did not stay on either activity or temperature pattern significantly longer (Fig. 4C, *P* = 0.1057).

### Tissue damage is weakly correlated to survival time and is driven by splenic damage

Tissue sections were stained with H&E, and a board-certified pathologist scored each tissue blindly for damage using a scale of 0 to 4 (0 = normal to 4 = severe damage, Supp. Table 1 and Supp. Fig. 4). Minimal damage was noted in the intestinal sections examined (ileum, cecum, and colon, scores 0-1) and these were excluded from further analyses (Supp. Fig. 5). To assess the relationship between survival and tissue damage, linear regression was performed showing survival and combined spleen and liver pathology scores with R^2^= 0.1256, while the spleen pathology score alone was 0.1323 and the liver pathology score alone was 0.08382 (Fig. 5B, C, D, all *P* = <0.001).

**Figure 5:**
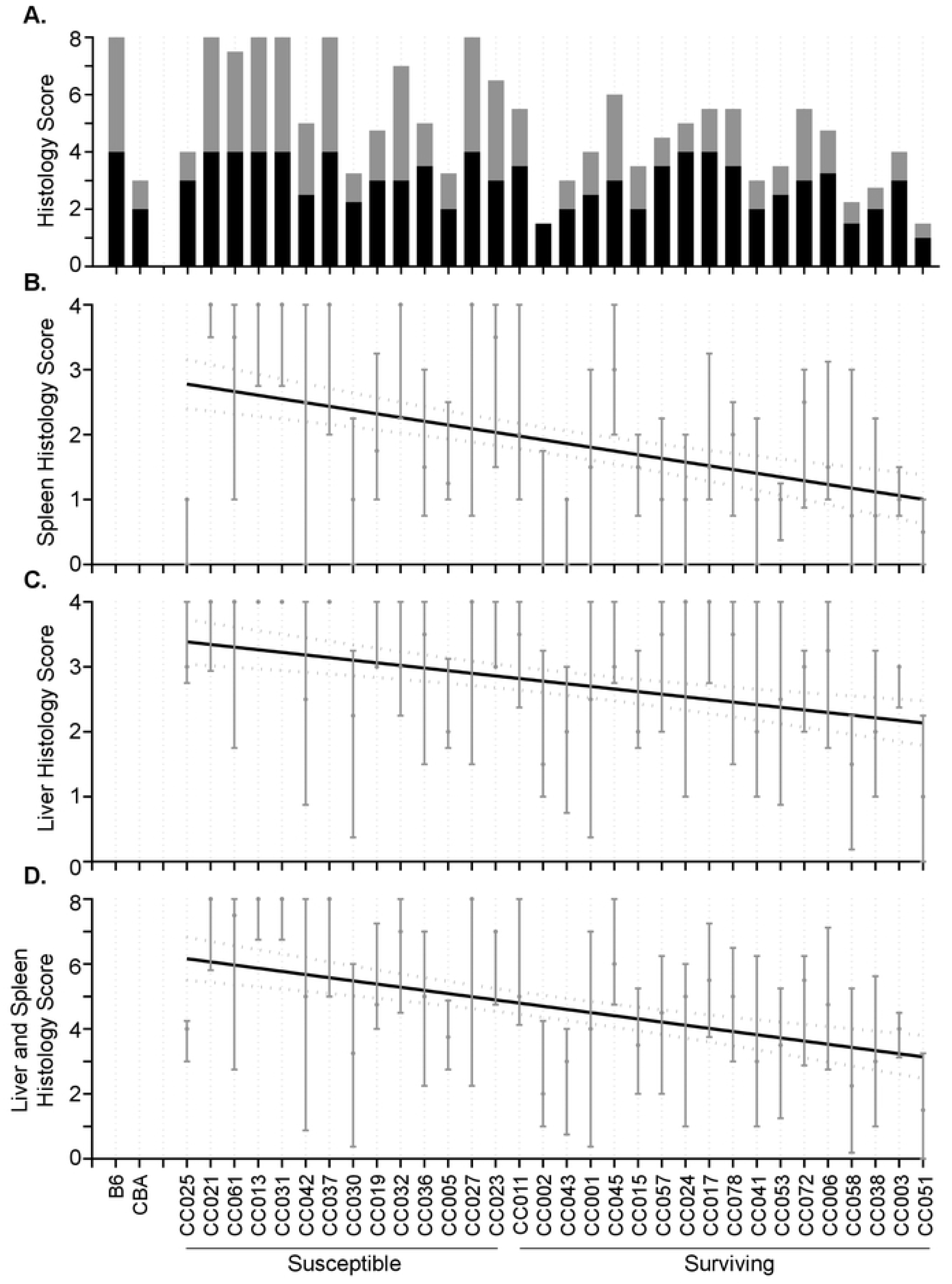
Spleen and liver damage correlate weakly with survival. A. Median Liver (black) and spleen (grey) histopathology scores for each strain. Scored on a scale of 0 to 4, with 0 being normal tissue and 4 being severely damaged tissue. B. Spleen (R^2^ = 0.1323, *P* = <0.0001) and C. liver (R^2^ = 0.08382, *P* = <0.0001) individual linear regression of histopathology scores. D. Linear regression of CC strains on combined spleen and liver histopathology scores (R^2^ = 0.1256, *P* = <0.0001). 95% confidence interval of regressions line is shown by dotted line. Medians and interquartile ranges are indicated.

Thus, damage to the spleen appears to be driving the small but significant correlation between histopathology scores and survival. Organs from susceptible strains were not necessarily more damaged than those from surviving strains, perhaps because the mice succumbed to infection before more severe damage could occur (Fig. 5A).

### Complete blood counts and circulating cytokines/chemokines do not explain differential host response to STm infection

Complete blood counts (CBC) were performed on pre-infection blood and on blood collected at necropsy (S1 Table and S2 Table). The differential counts for circulating white blood cell (WBC), neutrophils (NEU), monocytes (MON), lymphocytes (LYM) were analyzed. No significant differences were observed between surviving and non-surviving strains (Supp. Fig. 6, *P* = 0.4415, 0.2833, 0.9031, and 0.2505). The levels of 36 serum cytokines and chemokines were also determined from serum collected at necropsy and compared to those of uninfected control animals from each strain (S3 Table and S4 Table). Susceptibility or survival was not significantly correlated to the cytokine profile. The cytokine profile was influenced more by strain than infection status. Highly STm susceptible CC042 mice had high levels of almost every cytokine, suggesting that the mice were suffering a cytokine storm (S4 Table).

### Known polymorphisms do not always predict infection outcome

A point mutation in *Slc11a1* (also known as *Nramp1*, or *Ity*) is associated with susceptibility to lethal systemic STm infection (2,14,16). B6 is the only CC founder strain to carry this mutation, and B6 is susceptible to systemic infection with STm. Only 5 CC strains in the panel analyzed – CC021, CC031, CC042, CC045, and CC061 – carry the *Slc11a1* mutation (Fig. 2). All of these strains were highly colonized in liver and spleen, and, with the exception of one strain (CC045), succumbed to infection before day 7. CC045 had a high bacterial burden but survived to day 7 post-infection. This result suggests that CC045 has compensatory mechanisms to overcome STm susceptibility caused by the *Slc11a1* mutation.

Another mutation previously linked to STm susceptibility is in *Ncf2*, although it is not as detrimental to murine survival after *Salmonella* infection as the *Slc11a1* mutation (18,19). Three of the CC founders (PWK/PhJ, CAST/EiJ, NZO/ HlLtJ) carry the *Ncf2* mutation, and 12 of the CC strains in the current panel – CC013, CC015, CC023, CC025, CC030, CC032, CC036, CC038, CC045, CC058, CC072, CC078 – also carry the *Ncf2* mutation (Fig. 2). Six out of these 12 strains did not survive to day 7 post-infection (CC013, CC023, CC025, CC030, CC032, CC036), while the remaining 6 survived.

*Tlr4* also has a known point mutation linked to STm susceptibility, but none of the CC founder strains carry a mutation in this gene, making it impossible to verify the effect of this mutation on STm infection using CC mice (21,22).

Polymorphisms in *Ncf2* and *Slc11a1* combined explained STm susceptibility in 10 of the 14 susceptible strains, but the phenotypes of 4 of the strains – CC005, CC019, CC027, CC037 – are not explained by mutations in either of these genes. This finding suggests that other genetic factors contribute to STm susceptibility in these strains. CC045, the only strain to carry both *Slc11a1* and *Ncf2* mutations, is particularly unusual since 4 out of 6 mice of this strain survived STm infections.

### Genetic basis for variable response to infection with *Salmonella* Typhimurium

We used gQTL (46) to identify genetic regions associated with survival after oral STm infection. Since CC042 has a *de novo* mutation that explains its extreme susceptibility to STm infections, it was excluded from genetic analysis. CC045 was also excluded, as it carries mutations in both *Slc11a1* and *Ncf2*, yet survives infection. Using median survival time and percent of mice surviving to day 7 post-infection, significant associations were detected on Chromosomes (Chr) 4, 6, and 7 and suggestive associations were identified on Chr 1 and 2 (Fig. 6A and C, Table 1). We identified a significant association on Chr 4 (*Stq1* (Survival Time QTL)) and a significant association on Chr 7 (*Stq2*). The heritability of survival time was 45.55. Although the slope of survival percentages over time did not return significant temporal QTL, suggestive associations overlapped with those from the other survival parameters (Supp. Fig. 7). Other parameters such as weight change, CFUs, and CBCs were also analyzed using gQTL with no significant or suggestive associations detected, further supporting their independence from STm infection survival.

**Figure 6:**
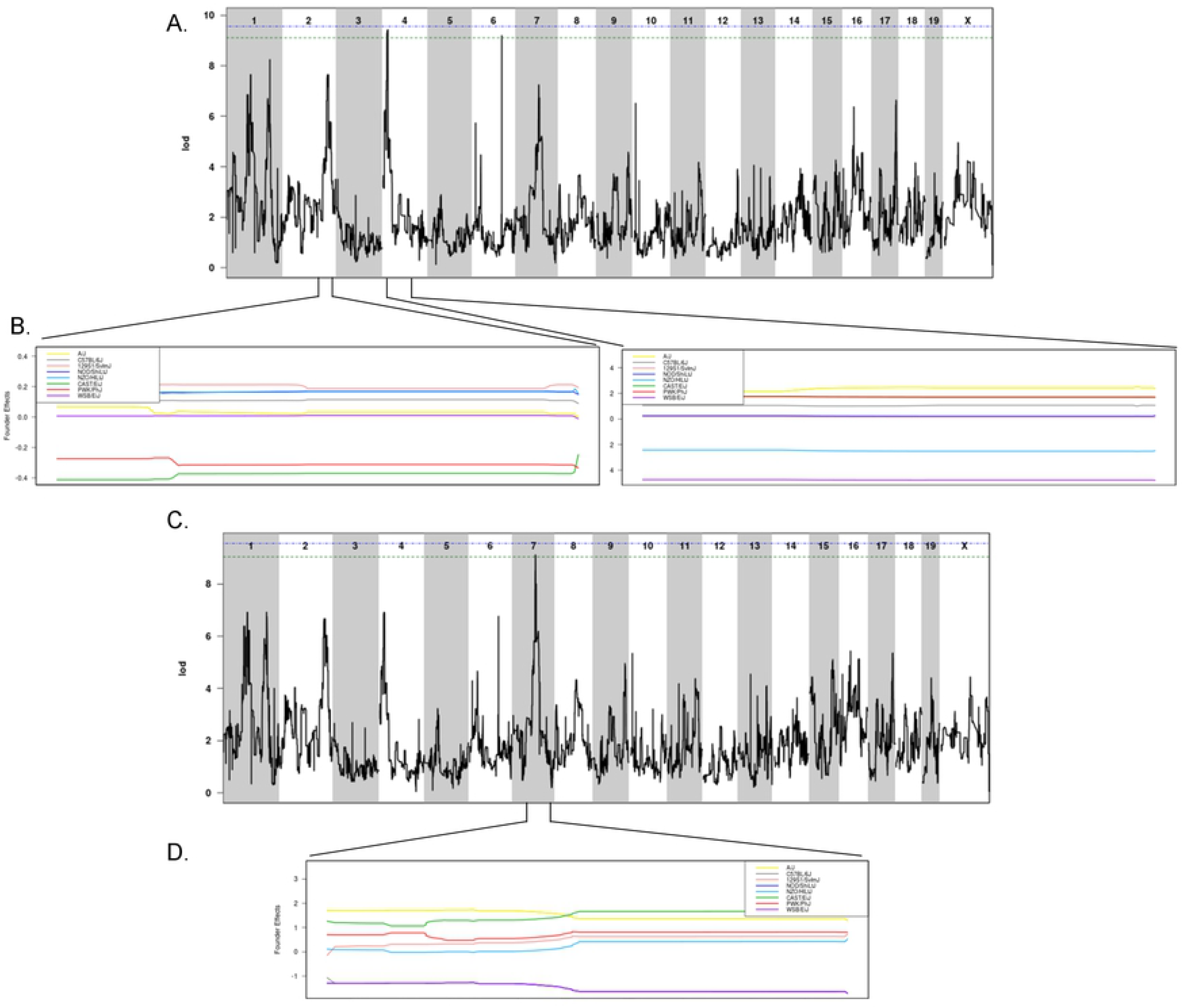
QTLs for survival and percent of mice surviving identified using 30 CC strains. A. QTL of median time survived after infection and B. allele effect plots focused on Chr 2 and Chr 4. C. QTL of percent of mice surviving to day 7 and D. allele effect plots focused on Chr 7. Green line designates 85% significance, blue line designates 90% significance. Results obtained using gQTL.

**Table 1:**
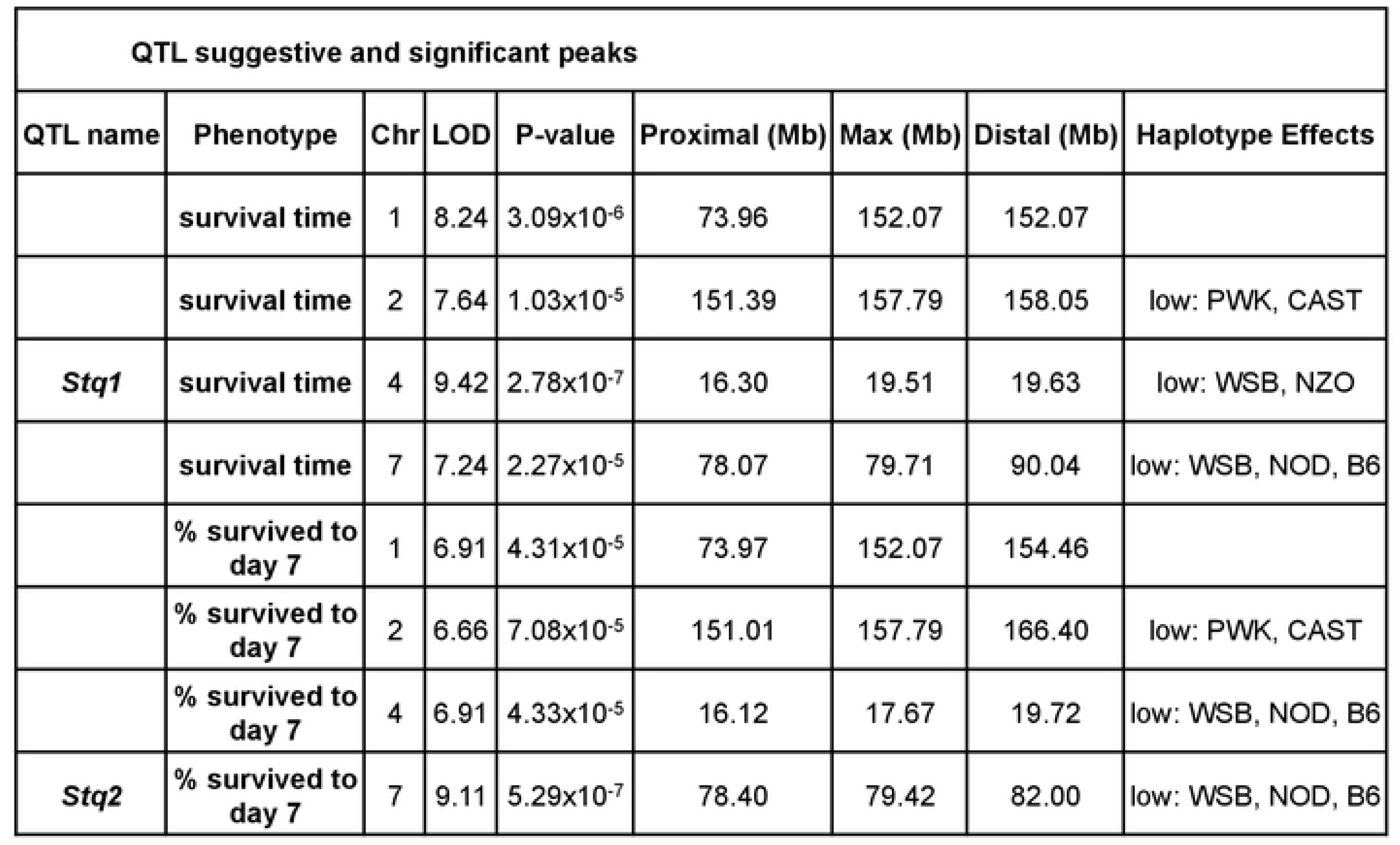
QTL associations for median survival time and percent survived to day 7 after STm infections. 30 CC strains were included in the gQTL analysis, excluding CC042 and CC045.

Two suggestive associations on Chr 1 overlapped with the *Slc11a1* and *Ncf2* genes, suggesting that these genes underlie these associations. The significant association on Chr 6 contains four genes, three hypothetical genes and one annotated gene, *Cntn3*, which is involved in nervous system development (Supp. Fig. 8A). However, no SNP differences were found in *Cntn3* that were consistent with founder haplotype effects. Although genes on Chr 2 and 4 have previously been shown to be involved in STm infections, the QTL regions we identified do not overlap with previously detected loci. The association on Chr 7 overlaps with a previously detected QTL region called *Ses2* (*S. enteritidis* susceptibility locus 1) which is linked to *Salmonella* clearance in the spleen, but the causative gene in this region remains unknown (47,48).

Using founder effect plots (Fig. 6 and Supp. Fig. 8 and 9), haplotype effects were determined for each QTL association. For median survival time, having a CC founder haplotype of PWK or CAST in the Chr 2 QTL region led to poor survival (Fig. 6B, Table 1). *Stq1* (Chr 4) was linked to poor survival if the region was inherited from the CC founders WSB and NZO (Fig. 6B, Table 1). B6, NOD, and WSB had low effect haplotypes for *Stq2* (Chr 7), leading to poor survival (Supp. Fig. 8B, Table 1). For percent of mice that survived to day 7, PWK and CAST alleles on the Chr 2 region led to poor survival, (Supp. Fig. 9A, Table 1), *Stq1* (Chr 4) was linked to poor survival for B6, WSB, and NOD alleles in this region (Supp. Fig. 9B, Table 1), and the *Stq2* (Chr 7) region was linked to poor survival for B6, NOD, and WSB (Fig. 6D, Table 1). No high haplotype effects or QTL linked to enhanced survival after STM infections were found in these associations.

To narrow the number of candidate genes for each association, SNP differences in coding regions in the CC founder strains were identified using the Mouse Phenome Database (The Jackson Laboratory). Ensembl Variant Effect Predictor (VEP) was used to determine whether the SNP difference had a predicted effect at the protein level. Haplotype effects for each locus were used to filter candidate genes that had differences in their sequences. For median survival time, Chr 2 had 259 genes with 13 SNP differences in genes that matched haplotype differences (Table 2). *Stq1*, with 23 genes, and Chr 7, with 232 genes, had no candidate genes with SNP differences. For percent of mice that survived to day 7, the Chr 2 region had 503 genes, of these genes 51 had SNP differences matching haplotype effects (Table 3). The Chr 4 region had 28 genes and *Stq2* had 115 genes, but neither region had SNP differences corresponding to haplotype differences.

**Table 2:**
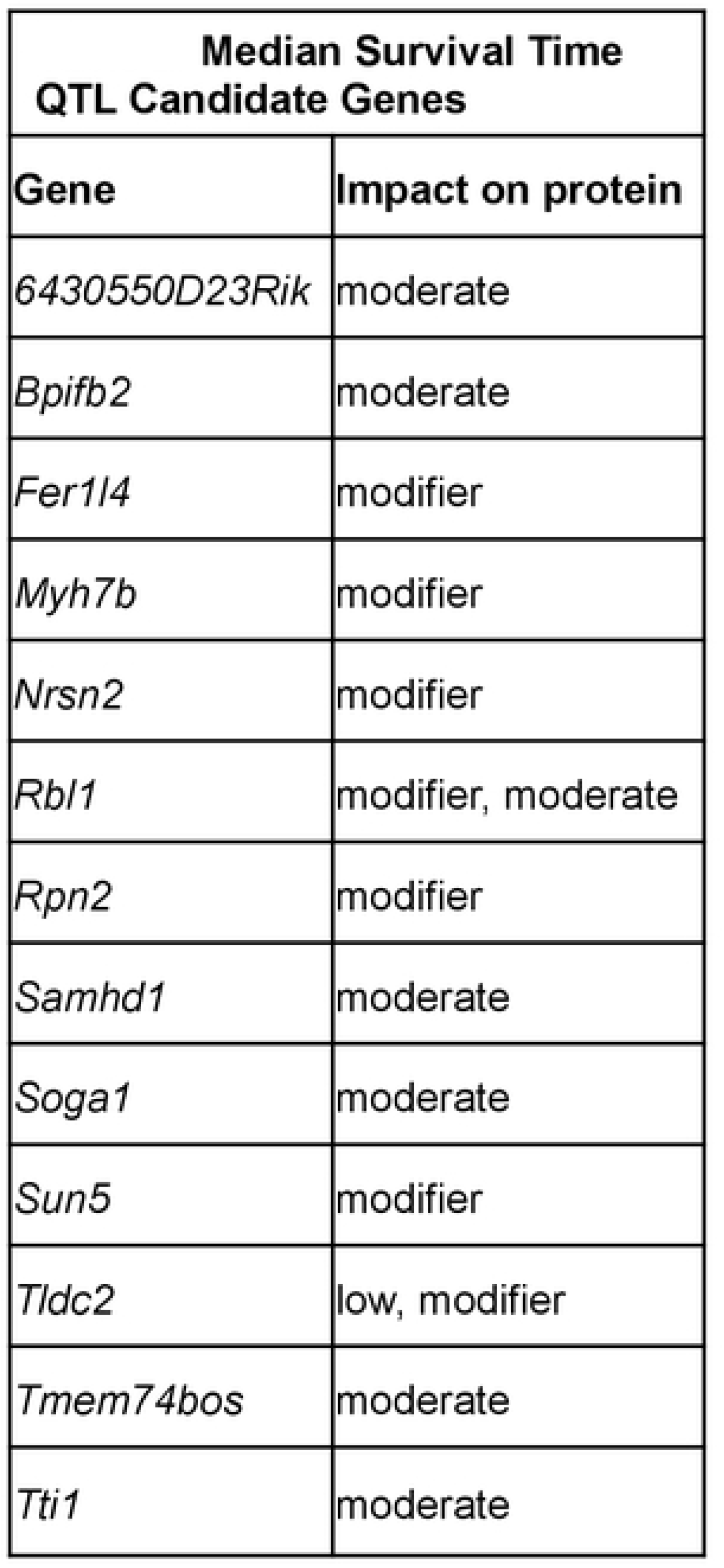
Candidate genes from median survival QTL associations that had SNP differences corresponding to haplotype differences. The impact the SNP difference is predicted to have on the resulting protein is shown.

**Table 3:**
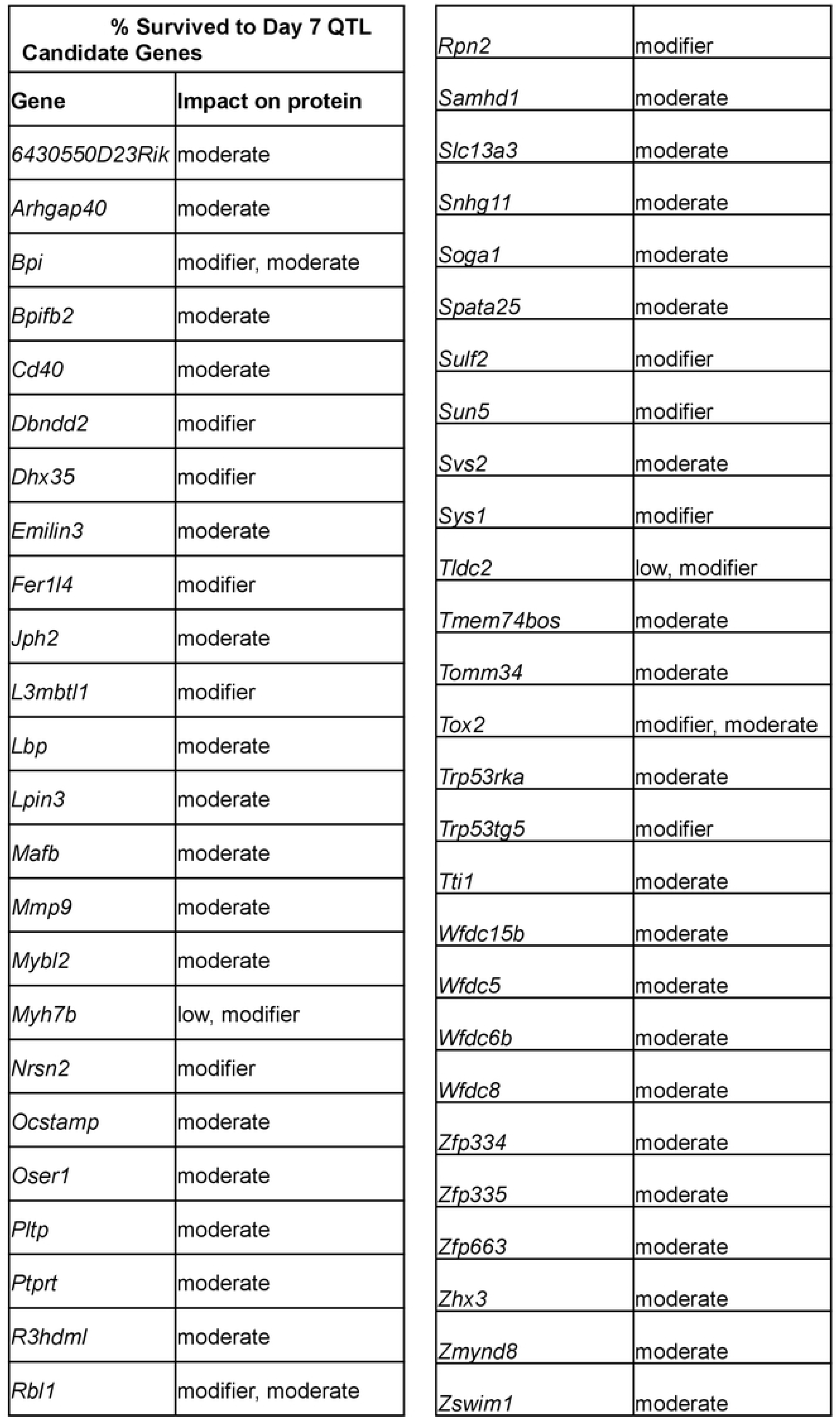
Candidate genes from percent survived to day 7 QTL associations that had SNP differences corresponding to haplotype differences. The impact the SNP difference is predicted to have on the resulting protein is shown.

## Discussion

Although CC strains have been extensively used to identify genes influencing susceptibility to viral infection (34–36,49–53), host determinants influencing the outcomes of bacterial infection have only been addressed in a small number of studies (37,39–41). We orally infected 32 CC strains of mice to study outcomes of STm infection in genetically diverse strains of mice. Our goal in this 7 day screen of acute STm infection was to identify the range of survival and colonization phenotypes and putative genetic differences behind these phenotypes.

STm colonization and survival of CC mice after oral STm infection were highly variable by strain, but were quite consistent within a strain. The 32 CC strains we tested differed in their range of colonization in spleen >20,000 fold (2.92 x10^2^ to 7.8 x10^8^ CFU/g) and >1 x 10^6^ fold (9.3 x10^1^ to 1.02 x10^8^ CFU/g) in liver. This wide range of colonization phenotypes reflects the genetic diversity of the CC, and is a wider range than represented in previous studies of the CC with STm (40,41). We identified 14 CC strains that were very susceptible to STm infection that did not survive the 7 day screening period. Several of these strains (CC025, CC021, CC013, and CC031) developed signs of systemic infection in the same time frame as B6 (susceptible control), but with lower bacterial colonization of both liver and spleen, suggesting increased sensitivity to STm infection. One of the susceptible strains, CC019, is also susceptible to intravenous infection with *M. tuberculosis* (37). Mice in the CC042 strain developed very high bacterial loads in liver and spleen in our studies, consistent with previous reports on intravenous infection with STm and infection with *M. tuberculosis* (40,41). The phenotype of CC042 mice has been linked to a mutation in the integrin alpha L gene (*Itgal*, also known as *Cd11a*) resulting in a defect in T cell recruitment (39,41). Finally, the remaining 18 CC strains we screened survived the full study period. Two of these strains, CC001 and CC002, were previously shown to display resistance to *M. tuberculosis* infection (37).

The range of survival for CC strains in our experiments was four to seven days (median), similar to those observed for B6 and CBA mice in our acute infection paradigm. The limited survival range is likely due to technical parameters of our experiment. Four days appears to be the fastest mice display symptoms of serious systemic disease after STm infection using our current infection paradigm. Seven days, the length of our screening period, is not long enough to capture the variability of longer survival after acute infection. Previous studies using CC mice and STm infection have not captured variability in survival after acute infection, because all mice were euthanized at day 4 post-infection regardless of disease state (40).

A few CC strains had unique infection profiles relative to those of traditional inbred mouse strains. CC051 mice were very poorly colonized and appeared healthier than CBA mice (resistant control). Thus, CC051 mice are more resistant to STm infection than previously studied strains, results consistent with low level colonization after intravenous STm infection (40). CC045 mice have mutations in both *Slc11a1* and *Ncf2*, both typically leading to poor survival after STm infection (14–16,18,19). While CC045 mice had higher bacterial loads of STm similar to susceptible mice, they survived the full study period and lost only ∼7% of their body weight. Thus, CC045 is a candidate “tolerant” strain, as it survives STm infection in the context of a bacterial colonization load that would be lethal to an otherwise susceptible mouse. CC045 mice must have compensatory mechanisms for survival despite the *Slc11a1* and *Ncf2* mutations that remain to be discovered.

Sex is well known to influence both innate and adaptive immune responses (54) as well as infection outcome (28,42,43,55,56). We identified several CC strains that exhibited sex differences in survival after STm infection (CC013, CC042, CC030, and CC027, Figure 2). In keeping with the general hypothesis that females have stronger innate and adaptive immune responses than males and are less susceptible to many infections (56), CC027 and CC042 females survived longer than males. CC027 exhibited the most dramatic sex difference as males were susceptible and all became moribund during the study period, while females survived. Consistent with the survival data, CC027 males were more heavily colonized in spleen and liver than females: (males: 4.80 x10^6^ and 1.63 x10^6^ CFU/g, females: 9.33 x10^5^ and 8.87 x10^4^ CFU/g, respectively). However, for the remaining two strains that exhibited sex differences in survival, CC013 and CC030, males survived longer than females. This finding is not without precedent however, as female mice lacking leucine-rich repeat kinase-2 protein (*Lrrk2*) are more susceptible to S. *Typhimurium* infection than males (57). Future work focused on the sex differences we describe should uncover additional genes linked to sexual dimorphism in susceptibility to bacterial infection.

Telemetry allows tracking of each mouse’s body temperature and activity minute by minute (58). These readings allowed us to define each mouse’s unique diurnal pattern of body temperature and activity, and to pinpoint the minute that each mouse started to develop disease. On average, a change in diurnal rhythm of temperature patterns were observable by telemetry 4,101 minutes (2.85 days) before infected mice developed clinical signs observable to the eye (Supp. Fig. 10), similar to previous work (58). Furthermore, our experiments mice that survived STm infection had a lower baseline core body temperature prior to infection, a novel and unexpected finding. As fever is generally thought to be disease fighting (59–62), we expected mice with higher baseline temperatures to be able to mount a quicker fever response. Although female B6 mice reportedly have a higher baseline body temperature than males, it is not known whether this difference influences susceptibility to infection (63). Finally, lower ambient temperature can lower the temperature and some parts of the body and impair antiviral immune responses (64), but the effect of lower core temperature on susceptibility to bacterial infections is unexplored.

When evaluating pre-infection baseline activity, surviving mice spent more time resting than susceptible mice during ‘active’ periods. Increased rest in surviving strains is unlikely to be the cause of lower core body temperature in these strains because it only occurred during the active period, while the temperature difference was maintained across all periods of the day. Furthermore, there was no significant difference in how rapidly surviving and susceptible infected mice deviated from their normal circadian pattern of activity after infection. However, susceptible mice deviated from their temperature pattern significantly earlier than their activity pattern, suggesting that core temperature is a more sensitive measurement to detect early disease.

Tissue damage had a small but significant correlation with survival time. Spleen and liver exhibited severe damage across most strains. Splenic damage drove the small correlation suggesting that liver damage, while more severe, is better tolerated than splenic damage. Interestingly, there were some strains that exhibited quite severe damage in both organs yet survived (CC011, CC017, CC045, CC072, and CC078) and some strains that had milder damage and did not survive (CC005, CC025, and CC030). While mice are typically categorized into resistant and susceptible, this variable damage data suggests that there is more than one way to respond to infection. Discovery of these phenotypes highlights the usefulness of the CC as a way to assay responses that may be influenced by genetic diversity.

While we studied the response to an oral STm infection, others have examined STm colonization after intravenous injection (40). Intravenous injection bypasses the intestine and defenses in this niche. Eighteen CC strains were shared between our study and previous work (40). When median colonization in liver and spleen was compared, the oral infection route (current infection) produced higher bacterial burdens and a wider range of colonization in general (liver: 10^4^ to 10^6^ CFU/g vs 10^3^ to 10^4^ CFU/g and spleen:10^3^ to 10^7^ CFU/g vs 10^4^ vs 10^5^ CFU/g, excluding CC042) (Fig. 2). Possible reasons for the differences in colonization levels include different route of delivery, different time to necropsy and bacterial strain differences (40). CC051 was an exception, with lower bacterial counts in the spleen after the oral infection versus IV infection. This finding suggests that CC051 mice are better at clearing intestinal STm, are better able to contain STm in the intestine, or are better at clearing STm prior to sustained systemic infection. CC051 is a model to explore true resistance to STm infection as it was the only strain that appeared to control systemic spread and systemic growth of STm after oral infection (1,65).

We analyzed all of the parameters we collected to identify putative linked genomic regions using gQTL (46). Only survival time yielded significant, 0.85 or higher, associations. Intervals on Chr 1 contained *Slc11a1* and *Ncf2*, known genes that have mutations linked to poor survival after STm infection (2,14–16,18–20). One strain, CC045, with *Slc11a1* and Ncf2 mutations survived the entire study period, as did six of twelve strains with *Ncf2* mutations. These strains must have alternative/compensatory mechanisms that allow them to overcome *Slc11a1* and *Ncf2* mutations. CC045 was highly colonized in both spleen and liver, making it a candidate for ‘tolerance’ to STm infection. The genetic basis for the ability of CC045 to survive STm infection in the face of these two mutations requires further exploration.

In addition to the intervals on Chr 1, suggestive and significant intervals on Chr 2, 4, and 7 were also identified by gQTL analysis. The intervals on Chr 2 and 4 do not contain any genes previously linked to STm infection, but the interval on Chr 7 overlaps with a previously discovered QTL region, *Ses2* (47,48). Genes linked to STm susceptibility underlying *Ses2* interval have not been identified. These regions were narrowed further as described above into a candidate gene list.

The suggestive QTL interval on Chr 2 had 13 candidate genes linked to median survival time after considering haplotypes (Table 2). The suggestive interval on Chr 2 for percent of mice surviving to day seven had 51 candidate genes remaining (Table 3). Promising genes in this region include: BPI fold containing family B, member 2 (*Bpifb2*), and SAM domain and HD domain, 1 (*Samhd1*). *Bpifb2* is a member of the lipid-transfer protein family and is predicted to have bactericidal and permeability-increasing properties, making it a promising candidate for further investigation (66). *Samhd1* regulates interferon pathways in response to viral infections through NF-KB (67,68). While *Samhd1* is linked to the immune response in viral infections, it may play a previously unknown role in immunity against bacterial pathogens. Further investigation is needed to prove such a link. In both median survival and percent survived to day seven QTL intervals for Chr 4 and 7, haplotype differences did not help us narrow the gene list. RNA expression analysis, currently underway, should identify candidate genes for these intervals.

We have carefully characterized the clinical course of STm infection using a population of genetically diverse Collaborative Cross mice and novel tools including telemetry analysis. We have identified novel responses to STm infection, sex differences in survival after STm infection, and determined that lower baseline body temperature is correlated with survival of STm infection. We defined CC strains bearing *Slc11a1* or *Ncf2* mutations that unexpectedly survive acute STm infection. Finally, we identified both known and novel regions of the mouse genome associated with survival time after STm infection. Overall, this research opens a rich area for discovery into the host genetics underlying the complex host response to bacterial infections.

## Acknowledgements

We thank Catherine Campbell, Yara Mohamed, Sandhiya Ravichandran, Kaya Mariello, and Connor Mathis for technical assistance. We also thank Dr. Phillip West for helpful discussions, and the Texas A&M Institute for Genome Sciences and Society (TIGSS) for use of their facility. This work was funded by the Defense Advanced Research Project Agency (DARPA), project DARPA D17AP00004.

## Materials and Methods

### Bacterial strains and media

The *Salmonella enterica* ser. Typhimurium strain (HA420) used for this study was derived from ATCC14028. HA420 is a fully virulent, spontaneous nalidixic acid resistant derivative of ATCC14028 (69). Strains were routinely cultured in Luria-Bertani (LB) broth and plates, supplemented with antibiotics when needed at 50 mg/L nalidixic acid (Nal).

For murine infections, strains were grown aerobically at 37°C to stationary phase in LB broth with nalidixic acid and diluted to generate an inoculum of 2-5 x10^7^ organisms in 100 microliters. Bacterial cultures used as inocula were serially diluted and plated to enumerate colony forming units (CFU) and determine the exact titer.

### Murine Strains

Both conventional mice (C57BL/6J and CBA/J) and Collaborative Cross (CC) mice were utilized in these experiments. All Collaborative Cross strains were obtained from UNC’s Systems Genetics Core Facility (SGCF) and either used directly for experiments or subsequently bred independently at Division of Comparative Medicine at Texas A&M University prior to these experiments (Supp. Table 2). Our experiments utilized 32 strains of CC mice, 3 females and 3 males per strain for a total of 228 mice (CC045 only had 2 females due to a surgical complication) (Supp. Table 2). Mice were fed Envigo Teklad Global 19% Protein Extruded Rodent Diet (2919) or Envigo Teklad Rodent Diet (8604) based on strain need.

### Ethics Statement

All animal experiments were conducted in accordance with the Guide for the Care and Use of Laboratory Animals, were reviewed by the TAMU IACUC and under AUP#s 2018-0488 D and 2015-0315 D.

### Placement of Telemetry Devices

5 to 9-week-old mice (C57BL/6, CBA, CC strains) were anesthetized with an isoflurane vaporizer for surgical telemetry device placement. The abdomen was opened with a midline abdominal incision (up to 2 cm). Starr life science G2 E-mitter devices were loosely sutured to the ventral abdominal wall. The abdominal muscle layer was closed with 5-0 vicryl, and the skin layer was closed using stainless steel wound clips (FST 9mm). Animals were given an intraperitoneal injection of Buprenorphine (0.0001 mg/g) for pain prior to recovery from anesthesia, and every 8 hours thereafter as necessary for pain control. After recovery from anesthesia, implanted mice were group-housed and monitored twice daily for pain and wound closure for 7 days post-surgery, when the clips were removed. Any animals found to have serious complications after surgery were humanely euthanized.

### Infection with *Salmonella* Typhimurium

After 4-7 days of acclimation in the BSL-2 facility, 8 to 12-week-old implanted mice were weighed and infected by gavage with a dose of 2-5 x 10^7^ CFU of *S.* Typhimurium HA420 in 100 microliters of LB broth. Infected mice were monitored twice daily for signs of disease and activity by visual inspection. When telemetry and health condition data suggested the development of clinical disease from infection, mice were humanely euthanized. Animals that remained clinically healthy throughout the duration of the experiment were humanely euthanized at 7 days post-infection.

### Bacterial load determination

Mice were humanely euthanized by CO_2_ asphyxiation, and the spleen, liver, ileum, cecum, and colon were collected. A third of each organ was collected in 3 mL of ice-cold PBS, weighed, homogenized, serially diluted in PBS, and plated on Nal plates for enumeration of S. Typhimurium in each organ. Data are expressed as CFU/g of tissue.

### Telemetry Monitoring

Prior to placing implanted mice on telemetry platforms, mice were moved into individual cages and provided with a cardboard hut and bedding material. Individual cages containing implanted mice were placed onto ER4000 receiver platforms, and body temperature (once per minute) and gross motor activity (continuous measurement summed each minute) data was collected. Body temperature and gross motor activity data were collected for 4-7 days pre-infection. Mice were removed briefly from the receiver platforms for infection and then were placed back on the platforms and data collection by telemetry was resumed. Infected animals were continuously monitored by telemetry in additional to twice daily visual assessment.

### Identification of deviation from circadian pattern of body temperature

Additional clinical information, such as the time of inoculation relative to the start of the experiment (denoted T), was reported and used in centering time series for comparison between mice (typically, seven days after the beginning of monitoring). Quantitative detection of deviation from the baseline “off-pattern,” using temperature data was calculated on an individual basis. A temperature time series was filtered, a definition of healthy variation was defined, then the time of first “off-pattern” was calculated using that definition on post-inoculation data.

Each mouse time series was preprocessed using a moving median filter with a one-day window. For a specific minute t, the median collection of temperature values from [t-720, t+720] was used in calculating a median for the value t. After this processing, healthy variation was defined as any temperature falling within the range of minimum to maximum values during the pre-inoculation phase [T-5760,T] (5760 minutes = 4 days). This choice allowed for enough data to account for natural inter-day variation due to potential factors such as inter-strain variation, epigenetic differences, and sex differences, while avoiding bias due to observed acclimation time after transfer to a new facility in some mice in the first few days of observation (Supp. Fig. 11A).

Identifying post-inoculation off-pattern behavior was done by identifying temperature values that fall outside the interval of healthy variation. The post-inoculation interval ranges from [T+60,T+10080] (7 days), where the one hour gap was used to avoid false positive detection due to the physical disturbance associated with inoculation (Supp. Fig. 11A).

### Detection of deviation from circadian pattern of activity

While activity data does exhibit circadian patterns, this data necessitated a different approach to preprocessing compared to temperature data because activity values are inherently non-negative and have a modal value of zero. Two approaches were taken. One was from the perspective of parameter estimation of iid sampling from a statistical distribution. The second approach was based on plainly calculating the fraction of non-zero activity values measured in a moving window. Hence, we approached the analysis of activity data from the perspective of determining the parameters of a stochastic process. In the first approach, for a given time interval t, we worked from the assumption that the number of activity values observed to be i obeys the following distribution:

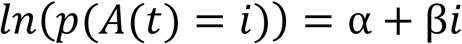

where the coefficient β, expected to be negative, corresponds to the modeling assumption that the relative drop in observed activity counts ought to decay (note 0<e^β<1 if β is negative). For instance, if the value of β= -0.693, so that e^β^ ≈1\/2, this would represent an assumption that there are half as many activity values observed to be 1 (one movement per minute) than 2 (two movements per minute). In actuality, this decay coefficient is typically seen to be β≈-0.025 to β≈-0.015, representing that observed activity values decay by half around every 27-46 values.

This theory is implemented in practice by windowing a mouse’s activity time series, creating a binning (empirical distribution) (*i*, *c*(*i*)),, the performing a log-linear fit by transforming *c*(*i*) → *ln* (1 + *c*(*i*)) and applying linear least-squares regression to calculate α and β. Statistical models with simpler assumptions stemming from a “memoryless” assumption were attempted but did not see any feasible agreement with observed activity data (Supp. Fig. 11B).

The second approach mentioned above produced a qualitatively cleaner signal, which again is based on windowing a mouse’s activity time series, and calculating the fraction of activity values in a window for which A(t)>0. One might expect not weighting by intensity of activity loses information, but we saw this approach to filtering activity data resulted in a time series for which we could apply the same methods as with the temperature time series (Supp. Fig 11B)

### Histopathology

After euthanasia, samples of liver, spleen, ileum, cecum, and colon from each mouse were collected and fixed in 10% neutral buffered formalin at room temperature for 24 hours and stored in 70% ethanol before embedding in paraffin, sectioning at 5 µm and staining with hematoxylin and eosin (H&E). Histologic sections of all tissues were evaluated in a blinded manner by brightfield microscopy and scored on a scale from 0 to 4 for tissue damage by a board-certified veterinary pathologist (Supp. Table 1 and Supp. Fig. 4). The combined scores of spleen and liver from individual mice were used to calculate the median and interquartile range for each group. Whole slide images of liver and spleen H&E-stained sections were captured as digital files by scanning at 40x using a 3DHistech Pannoramic SCAN II FL™ scanner (Epredia, Kalamazoo, MI). Digital files were processed by Aiforia Hub™ (Aiforia, Cambridge, MA) software for generating the images with 100 µm scale bars.

### Complete Blood Count

For some mice, whole blood was collected by submandibular bleed for complete blood count (CBC) one week prior to surgery for the pre-infection sample to determine each mouse’s baseline. Blood was also collected at necropsy by cardiac puncture. Additional blood was collected from uninfected control animals by cardiac puncture after euthanasia and median values were used as the baseline for mice that were not their own pre-infection control. All blood was collected into EDTA tubes and analyzed on an Abaxis VetScan HM5.

### Cytokine and Chemokine Analysis

Serum was stored at -80°C and thawed on ice immediately prior to cytokine assays. Serum cytokine levels were evaluated using an Invitrogen ProcaratPlex Cytokine/Chemokine Convenience Panel 1A 36 plex kit as per manufacturer instructions (ThermoFisher). Briefly, magnetic beads were added to each well of the 96-well plate and washed on the Bio-Plex Pro Wash Station. 25uL of samples and standards were then added to the plate along with 25uL of universal assay buffer. Plates were then shaken at room temperature for 1 hour and washed. 25uL of detection antibody was added an incubated for 30 minutes and washed away, followed by 50uL of SAPE for 30 minutes and washed away. 120uL of reading buffer was added and plates were evaluated using a Bio-Plex 200 (BioRad). Samples and standards were run in duplicate, and samples were diluted as needed to get 25ul of serum per duplicate. The kit screens for the following 36 cytokines and chemokines: IFN gamma; IL-12p70; IL-13; IL-1 beta; IL-2; IL-4; IL-5; IL-6; TNF alpha; GM-CSF; IL-18; IL-10; IL-17A; IL-22; IL-23; IL-27; IL-9; GRO alpha; IP-10; MCP-1; MCP-3; MIP-1 alpha; MIP-1 beta; MIP-2; RANTES; Eotaxin; IFN alpha; IL-15/IL-15R; IL-28; IL-31; IL-1 alpha; IL-3; G-CSF; LIF; ENA-78/CXCL5; M-CSF. Median values of uninfected animals for each strain were used as the baseline for that strain.

### QTL analysis

gQTL, an online resource designed specifically to run CC QTLs, was used to identify putative QTLs (46). Briefly, median values of each strain for various parameters were uploaded to the website and QTLs were run using 1000 permutations with “automatic” transformation. Automatic picks either log or square root transformations, whichever normalizes the data best.

---

**Supplemental Figure 1: Survival is correlated to CFU and strongly correlated to % weight change.** Strain medians for 32 CC strains were analyzed using Spearman correlation. Blue detonates positive correlations and red detonates negative correlations.

**Supplemental Figure 2: Survival for susceptible strains is weakly correlated to CFU and strongly correlated to % weight change.** Strain medians for 14 CC strains that are susceptible were analyzed using Spearman correlation. Blue detonates positive correlations and red detonates negative correlations.

**Supplemental Figure 3: Each strain has a unique circadian pattern before infection.** Purple lines represent the average temperature and black lines represent the average fraction of time the mouse is active pre-infection for 7 days. The gray area represents +/- 2 standard deviations. A-N are susceptible strains, O-AF are surviving strains, and AG-AH are control strains.

**Supplemental Table 1: Scoring matrix with descriptions**. Final score per organ corresponds to the highest score assigned to a single category. Scoring matrix from Dr. L. Garry Adams.

**Supplemental Figure 4: Representative images of spleen and liver histology scoring scheme.** Tissues were sectioned and stained with H&E before being analyzed. Images from Dr. L. Garry Adams illustrating scoring matrix in Supp. Table 1.

**Supplemental Figure 5: Intestinal organs were minimally damaged after STm infections.** Median ileum (gray) and cecum/colon (black) histopathology scores. Scored on a logarithmic scale of 0 to 4, with 0 being normal tissue and 4 being severely damaged tissue.

**Supplemental Figure 6**: **White blood cell numbers are not significantly different between surviving and susceptible strains.** Strain medians for change in blood count (infected-uninfected) for A. total white blood cells, B. neutrophils, C. monocytes, and D. lymphocytes, grouped by survival status.

**Supplemental Figure 7**: **QTL of slope of survival curve.** This value defines how rapidly each strain succumbed to infection. QTL analysis included 30 CC strains, excluding CC042 and CC045.

**Supplemental Figure 8: Founder effects plots for QTL of median time survived**. 30 CC strains, included, after removal of CC042 and CC045. Founder effect plots for A. Chr 6 and B. 7 focused on the QTL peak, genes within the region are also shown. Results obtained using gQTL analysis.

**Supplemental Figure 9: Founder effect plots of QTL of percent of mice surviving to day 7.** Analysis included 30 CC strains, after removal of CC042 and CC045. Founder effect plots for A. Chr 2 and B. 4 focused on the QTL peak, genes within the region are also shown. Results obtained using gQTL analysis.

**Supplemental Figure 10: 32 CC strains have earlier disruption of circadian pattern than clinical signs of disease.** Time from first change in temperature pattern subtracted from time that mouse developed visually observable clinical signs of disease. Mean and standard deviation shown by strains and individual mice shown by dots.

**Supplemental Table 2: Number of mice used and facility of origin for 2 control strains and 32 CC strains.**

**Supplemental Figure 11**: **Telemetry calculations images.** Examples of how A. temperature and B. activity calculations were determined. Specifically maximum, median, minimum pre-infection temperature and activity as well “off-pattern” for post-infection.

**S1 Table: Baseline Complete Blood Counts from uninfected mice for 32 CC strains and 2 control strains (B6 and CBA).** Mean and standard deviation are shown. See excel sheet.

**S2 Table: Change in Complete Blood Counts pre- to post-infection.** The difference between white blood cell counts after infection versus pre-infection was determined for 32 CC strains and 2 control strains (B6 and CBA). Mean and standard deviation are shown. See excel sheet.

**S3 Table: Baseline level for 36 cytokines prior to infection.** Mean and standard deviation are shown for 28 CC strains as well as two controls (B6 and CBA). See excel sheet.

**S4 Table: Change in cytokine amount for each CC strain across infection.** The differences between cytokine amounts from the serum of infected animals and the cytokine amount from uninfected animals was determined for 28 CC strains and 2 control strains (B6 and CBA). Mean and standard deviation are shown. See excel sheet.

## References

1. Santos RL, Tükel Ç, Raffatellu M, Bäumler AJ, Tsolis RM, Adams LG, et al. Life in the inflamed intestine, Salmonella style. Trends Microbiol. 2009;17(11):498–506.

2. Gilchrist JJ, MacLennan C a., Hill AVS. Genetic susceptibility to invasive Salmonella disease. Nat Rev Immunol [Internet]. 2015;15(7):452–63. Available from: http://www.ncbi.nlm.nih.gov/pubmed/26109132

3. LaRock DL, Chaudhary A, Miller SI. Salmonellae interactions with host processes. Nat Rev Microbiol. 2015;13(4):191–205.

4. Majowicz SE, Musto J, Scallan E, Angulo FJ, Kirk M, O’Brien SJ, et al. The global burden of nontyphoidal salmonella gastroenteritis. Clin Infect Dis. 2010;50(6):882–9.

5. Collin R, Balmer L, Morahan G, Lesage S. Common Heritable Immunological Variations Revealed in Genetically Diverse Inbred Mouse Strains of the Collaborative Cross. J Immunol. 2019;202(3):777–86.

6. Stanaway JD, Parisi A, Sarkar K, Blacker BF, Reiner RC, Hay SI, et al. The global burden of non-typhoidal salmonella invasive disease: a systematic analysis for the Global Burden of Disease Study 2017. Lancet Infect Dis. 2019;19(12):1312–24.

7. CDC. Antibiotic resistance threats in the United States. Centers Dis Control Prev [Internet]. 2019;1–113. Available from: https://www.cdc.gov/drugresistance/biggest_threats.html

8. Nathan C. Antibiotics at the crossroads. Nature [Internet]. 2004;431(7011):899–902. Available from: http://www.nature.com/doifinder/10.1038/431899a

9. Threlfall EJ. Antimicrobial drug resistance inSalmonella: problems andperspectives in food- and water-borne infections. FEMS Microbiol Rev Elsivier Sci [Internet]. 2002;26:141–8. Available from: https://onlinelibrary.wiley.com/doi/epdf/10.1111/j.1574-6976.2002.tb00606.x

10. Patin E, Hasan M, Bergstedt J, Rouilly V, Libri V, Urrutia A, et al. Natural variation in the parameters of innate immune cells is preferentially driven by genetic factors resource. Nat Immunol. 2018;19(3):302–14.

11. Roy MF, Malo D. Genetic regulation of host responses to Salmonella infection in mice. Genes Immun. 2002;3(7):381–93.

12. de Jong HK, Parry CM, van der Poll T, Wiersinga WJ. Host-Pathogen Interaction in Invasive Salmonellosis. PLoS Pathog. 2012;8(10):1–9.

13. Tsolis RM, Xavier MN, Santos RL, Bäumler AJ. How to become a top model: Impact of animal experimentation on human Salmonella disease research. Infect Immun. 2011;79(5):1806–14.

14. Lalmanach AC, Montagne A, Menanteau P, Lantier F. Effect of the mouse Nramp1 genotype on the expression of IFN-γ gene in early response to Salmonella infection. Microbes Infect. 2001;3(8):639–44.

15. Gopinath S, Hotson A, Johns J, Nolan G, Monack D. The Systemic Immune State of Super-shedder Mice Is Characterized by a Unique Neutrophil-dependent Blunting of TH1 Responses. PLoS Pathog. 2013;9(6).

16. Mastroeni P, Ugrinovic S, Chandra A, MacLennan C, Doffinger R, Kumararatne D. Resistance and susceptibility to Salmonella infections: Lessons from mice and patients with immunodeficiencies. Rev Med Microbiol. 2003;14(2):53–62.

17. Govoni G, Gros P. Macrophage NRAMP1 and its role in resistance to microbial infections. Inflamm Res. 1998;47(7):277–84.

18. Sancho-Shimizu V, Malo D. Sequencing, Expression, and Functional Analyses Support the Candidacy of Ncf2 in Susceptibility to Salmonella Typhimurium Infection in Wild-Derived Mice. J Immunol. 2006;176(11):6954–61.

19. Khan RT, Chevenon M, Yuki KE, Malo D. Genetic dissection of the Ity3 locus identifies a role for Ncf2 co-expression modules and suggests Selp as a candidate gene underlying the Ity3.2 locus. Front Immunol. 2014;5(AUG):1–13.

20. Khan RT, Yuki KE, Malo D. Fine-mapping and phenotypic analysis of the Ity3 salmonella susceptibility locus identify a complex genetic structure. PLoS One. 2014;9(1).

21. Li Q, Cherayil BJ. Role of toll-like receptor 4 in macrophage activation and tolerance during Salmonella enterica serovar Typhimurium infection. Infect Immun. 2003;71(9):4873–82.

22. Bihl F, Larivière L, Qureshi ST, Flaherty L, Malo D. LPS-hyporesponsiveness of mnd mice is associated with a mutation in Toll-like receptor 4. Genes Immun. 2001;2(1):56–9.

23. Welsh CE, Miller DR, Manly KF, Wang J, McMillan L, Morahan G, et al. Status and access to the Collaborative Cross population. Mamm Genome. 2012;23(9– 10):706–12.

24. Threadgill DW, Churchill GA. Ten years of the collaborative cross. Genetics. 2012;190(2):291–4.

25. Aylor DL, Valdar W, Foulds-mathes W, Buus RJ, Verdugo R a, Baric RS, et al. Genetic analysis of complex traits in the emerging Collaborative Genetic analysis of complex traits in the emerging Collaborative Cross. Genome Res. 2011;21(Churchill 2007):1213–22.

26. Churchill G, Airey D, Allayee H, … The Collaborative Cross, a community resource for the genetic analysis of complex traits. Nat Rev Genet. 2004;36(11):1133–7.

27. Philippi J, Xie Y, Miller DR, Bell TA, Zhang Z, Lenarcic AB, et al. Using the emerging Collaborative Cross to probe the immune system. Genes Immun. 2014;15(1):38–46.

28. Abu Toamih Atamni H, Nashef A, Iraqi FA. The Collaborative Cross mouse model for dissecting genetic susceptibility to infectious diseases. Mamm Genome [Internet]. 2018;29:471–87. Available from: http://link.springer.com/10.1007/s00335-018-9768-1

29. Smith CM, Sassetti CM. Modeling Diversity: Do Homogeneous Laboratory Strains Limit Discovery? Trends Microbiol [Internet]. 2018;26(11):892–5. Available from: https://www.cell.com/trends/microbiology/fulltext/S0966-842X(18)30174-4

30. Raberg L, Sim D, Read AF. Disentangling Genetic Variation for Resistance and Tolerance to Infectious Disease in Animals. Science (80- ). 2007;318(November):812–4.

31. Raberg L, Graham AL, Read AF. Decomposing health: tolerance and resistance to parasites in animals. Philos Trans R Soc B Biol Sci [Internet]. 2009;364(1513):37–49. Available from: http://rstb.royalsocietypublishing.org/cgi/doi/10.1098/rstb.2008.0184

32. Medzhitov R, Schneider DS, Soares MP, Caldwell RM, Schafer JF, Compton LE, et al. Disease tolerance as a defense strategy. Science [Internet]. 2012;335(6071):936–41. Available from: http://www.ncbi.nlm.nih.gov/pubmed/22363001%5Cnhttp://www.pubmedcentral.nih.gov/articlerender.fcgi?artid=PMC3564547

33. Maurizio PL, Ferris MT, Keele GR, Miller DR, Shaw GD, Whitmore AC, et al. Bayesian Diallel Analysis Reveals *Mx1* -Dependent and *Mx1* -Independent Effects on Response to Influenza A Virus in Mice. G3: Genes|Genomes|Genetics [Internet]. 2017;8(February):g3.300438.2017. Available from: http://g3journal.org/lookup/doi/10.1534/g3.117.300438

34. Rasmussen AL, Okumura A, Ferris MT, Green R, Feldmann F, Kelly SM, et al. Host genetic diversity enables Ebola hemorrhagic fever pathogenesis and resistance. Science (80- ). 2014;346(6212):987–91.

35. Kollmus H, Pilzner C, Leist SR, Heise M, Geffers R, Schughart K. Of mice and men: the host response to influenza virus infection. Mamm Genome [Internet]. 2018;0(0):1–25. Available from: http://dx.doi.org/10.1007/s00335-018-9750-y

36. Brinkmeyer-Langford CL, Rech R, Amstalden K, Kochan KJ, Hillhouse AE, Young C, et al. Host genetic background influences diverse neurological responses to viral infection in mice. Sci Rep [Internet]. 2017;7(1):1–17. Available from: http://dx.doi.org/10.1038/s41598-017-12477-2

37. Smith CM, Proulx MK, Olive AJ, Laddy D, Mishra BB, Moss C, et al. Tuberculosis susceptibility and vaccine protection are independently controlled by host genotype. MBio. 2016;7(5):1–13.

38. Green R, Wilkins C, Thomas S, Sekine A, Ireton RC, Ferris MT, et al. Identifying protective host gene expression signatures within the spleen during West Nile virus infection in the collaborative cross model. Genomics Data [Internet]. 2016;10:114–7. Available from: http://linkinghub.elsevier.com/retrieve/pii/S2213596016301386

39. Smith CM, Proulx MK, Villena FP De, Lai R, Kiritsy MC, Bell TA, et al. Functionally Overlapping Variants Control Tuberculosis Susceptibility in Collaborative Cross Mice. MBio. 2019;10(6):1–15.

40. Zhang J, Malo D, Mott R, Panthier J, Montagutelli X, Jaubert J. Identification of new loci involved in the host susceptibility to Salmonella Typhimurium in collaborative cross mice. BMC Genomics. 2018;19(303):1–13.

41. Zhang J, Teh M, Kim J, Eva MM, Cayrol R, Meade R, et al. A Loss-of-Function Mutation in the Integrin Alpha L (Itgal) Gene Contributes to Susceptibility to Salmonella enterica Serovar Typhimurium Infection in Collaborative Cross Strain CC042. Infect Immun. 2019;88(1):1–19.

42. Geurs TL, Hill EB, Lippold DM, French AR. Sex Differences in Murine Susceptibility to Systemic Viral Infections. J Autoimmun [Internet]. 2012;38(2–3). Available from: https://www.ncbi.nlm.nih.gov/pmc/articles/PMC3624763/pdf/nihms412728.pdf

43. Boivin GA, Pothlichet J, Skamene E, Brown EG, Loredo-Osti JC, Sladek R, et al. Mapping of Clinical and Expression Quantitative Trait Loci in a Sex-Dependent Effect of Host Susceptibility to Mouse-Adapted Influenza H3N2/HK/1/68. J Immunol. 2012;188(8):3949–60.

44. Failla KR, Connelly CD. Systematic Review of Gender Differences in Sepsis Management and Outcomes. J Nurs Scholarsh. 2017;312–24.

45. Aminian M, Andrews-Polymenis H, Gupta J, Kirby M, Kvinge H, Ma X, et al. Mathematical methods for visualization and anomaly detection in telemetry datasets. Interface Focus. 2020;10(1).

46. Konganti K, Ehrlich A, Rusyn I, Threadgill D. gQTL : A Web Application for QTL Analysis Using the Collaborative Cross Mouse Genetic Reference Population. Genes Genomes Genet. 2018;8(10).

47. Caron J, Loredo-Osti JC, Laroche L, Skamene E, Morgan K, Malo D. Identification of genetic loci controlling bacterial clearance in experimental Salmonella enteritidis infection: An unexpected role of Nramp1 (Slc11a1) in the persistence of infection in mice. Genes Immun. 2002;3(4):196–204.

48. Caron J, Loredo-Osti JC, Morgan K, Malo D. Mapping of interactions and mouse congenic strains identified novel epistatic QTLs controlling the persistence of Salmonella Enteritidis in mice. Genes Immun. 2005;6(6):500–8.

49. Graham JB, Thomas S, Swarts J, McMillan AA, Ferris MT, Suthar MS, et al. Genetic diversity in the collaborative cross model recapitulates human west nile virus disease outcomes. MBio. 2015;6(3):1–11.

50. Graham JB, Swarts JL, Wilkins C, Thomas S, Green R, Sekine A, et al. A Mouse Model of Chronic West Nile Virus Disease. PLoS Pathog. 2016;12(11):1–23.

51. Ferris MT, Aylor DL, Bottomly D, Whitmore AC, Aicher LD, Bell TA, et al. Modeling Host Genetic Regulation of Influenza Pathogenesis in the Collaborative Cross. PLoS Pathog. 2013;9(2).

52. Price A, Okumura A, Haddock E, Feldmann F, Meade-White K, Sharma P, et al. Transcriptional Correlates of Tolerance and Lethality in Mice Predict Ebola Virus Disease Patient Outcomes. Cell Rep. 2020;30:1702–13.

53. Gralinski LE, Ferris MT, Aylor DL, Whitmore AC, Green R, Frieman MB, et al. Genome Wide Identification of SARS-CoV Susceptibility Loci Using the Collaborative Cross. PLoS Genet. 2015;11(10):1–21.

54. Klein SL, Flanagan KL. Sex differences in immune responses. Nat Rev Immunol. 2016;16(10):626–38.

55. Zychlinsky Scharff A, Rousseau M, Lacerda Mariano L, Canton T, Consiglio CR, Albert ML, et al. Sex differences in IL-17 contribute to chronicity in male versus female urinary tract infection. JCI insight. 2019;5(13):1–19.

56. Jaillon S, Berthenet K, Garlanda C. Sexual Dimorphism in Innate Immunity. Clin Rev Allergy Immunol. 2019;56(3):308–21.

57. Shutinoski B, Hakimi M, Harmsen IE, Lunn M, Rocha J, Lengacher N, et al. Lrrk2 alleles modulate inflammation during microbial infection of mice in a sex-dependent manner. Sci Transl Med. 2019;11(511):36–41.

58. Williamson ED, Savage VL, Lingard B, Russell P, Scott EAM. A biocompatible microdevice for core body temperature monitoring in the early diagnosis of infectious disease. Biomed Microdevices. 2007;9(1):51–60.

59. Plaza JJG, Hulak N, Zhumadilov Z, Akilzhanova A. Fever as an important resource for infectious diseases research. Intractable Rare Dis Res. 2016;5(2):97–102.

60. Schieber AMP, Ayres JS. Thermoregulation as a disease tolerance as a defense strategy. Pathog Dis. 2016;74(9).

61. Vlach KD, Boles JW, Stiles BG. Telemetric evaluation of body temperature and physical activity as predictors of mortality in a murine model of staphylococcal enterotoxic shock. Comp Med [Internet]. 2000;50(2):160–6. Available from: http://www.ingentaconnect.com/content/aalas/cm/2000/00000050/00000002/art00010?token=00511e21ef0b2517308d39437a63736a6f3547634c3e666c24452a566f644a467b4d616d3f4e4b341

62. Elhadad D, McClelland M, Rahav G, Gal-Mor O. Feverlike temperature is a virulence regulatory cue controlling the motility and host cell entry of typhoidal Salmonella. J Infect Dis. 2015;212(1):147–56.

63. Sanchez-Alavez M, Alboni S, Conti B. Sex- and age-specific differences in core body temperature of C57Bl/6 mice. Age (Omaha). 2011;33(1):89–99.

64. Foxman EF, Storer JA, Fitzgerald ME, Wasik BR, Hou L, Zhao H, et al. Temperature-dependent innate defense against the common cold virus limits viral replication at warm temperature in mouse airway cells. Proc Natl Acad Sci U S A. 2015;112(3):827–32.

65. McLaughlin PA, Bettke JA, Tam JW, Leeds J, Bliska JB, Butler BP, et al. Inflammatory monocytes provide a niche for Salmonella expansion in the lumen of the inflamed intestine. PLoS Pathog [Internet]. 2019;15(7):1–17. Available from: http://dx.doi.org/10.1371/journal.ppat.1007847

66. Bingle CD, Bingle L, Craven CJ. Distant cousins: Genomic and sequence diversity within the BPI fold-containing (BPIF)/PLUNC protein family. Biochem Soc Trans. 2011;39(4):961–5.

67. Chen S, Bonifati S, Qin Z, Gelais CS, Kodigepalli KM, Barrett BS, et al. SAMHD1 suppresses innate immune responses to viral infections and inflammatory stimuli by inhibiting the NF-κB and interferon pathways. Proc Natl Acad Sci U S A. 2018;115(16):E3798–807.

68. Li Z, Huan C, Wang H, Liu Y, Liu X, Su X, et al. TRIM 21-mediated proteasomal degradation of SAMHD 1 regulates its antiviral activity. EMBO Rep. 2020;21(1):1–18.

69. Bogomolnaya LM, Santiviago CA, Yang HJ, Baumler AJ, Andrews-Polymenis HL. “Form variation” of the O12 antigen is critical for persistence of Salmonella Typhimurium in the murine intestine. Mol Microbiol. 2008;70(5):1105–19.

